# A primer genetic toolkit for exploring mitochondrial biology and disease using zebrafish

**DOI:** 10.1101/542084

**Authors:** Ankit Sabharwal, Jarryd M. Campbell, Zachary WareJoncas, Mark Wishman, Hirotaka Ata, Wiebin Liu, Noriko Ichino, Jake D. Bergren, Mark D. Urban, Rhianna Urban, Tanya L. Poshusta, Yonghe Ding, Xiaolei Xu, Karl J. Clark, Stephen C. Ekker

## Abstract

Mitochondria are a dynamic eukaryotic innovation that play diverse roles in biology and disease. The mitochondrial genome is remarkably conserved in all vertebrates, encoding the same 37 gene set and overall genomic structure ranging from 16,596 base pairs (bp) in the teleost zebrafish (*Danio rerio*) to 16,569 bp in humans. Mitochondrial disorders are amongst the most prevalent inherited diseases affecting roughly 1 in every 5000 individuals. Currently, few effective treatments exist for those with mitochondrial ailments, representing a major unmet patient need. Mitochondrial dysfunction is also implicated to be a common component of a wide variety of other human illnesses ranging from neurodegenerative disorders like Huntington’s disease and Parkinson’s disease to autoimmune illnesses such as multiple sclerosis and rheumatoid arthritis. The electron transport chain (ETC) component of mitochondria is critical for mitochondrial biology and defects can lead to many mitochondrial disease symptoms. Here we present a publicly available collection of genetic mutants created in highly conserved, nuclear-encoded mitochondrial genes in *Danio rerio*. The zebrafish system represents a potentially powerful new opportunity for the study of mitochondrial biology and disease due to the large number of orthologous genes shared with humans and the many advanced features of this model system from genetics to imaging. This collection includes 22 mutant lines in 18 different genes created by locus-specific gene editing to induce frameshift or splice acceptor mutations leading to predicted protein truncation during translation. Also included are 6 lines created by the random insertion of the gene-breaking transposon (GBT) protein trap cassette. All of these targeted mutant alleles truncate conserved domains of genes critical to the proper function of the ETC or genes that have been implicated in human mitochondrial disease. This collection is designed to accelerate the use of zebrafish to study of many different aspects of mitochondrial function with the goal of widening our understanding of their role in biology and human disease.

## Introduction

Mitochondria are semi-autonomous organelles critical for eukaryotic cell function. The mitochondrial endosymbiotic genesis origin hypothesis proposes its evolution from an alpha proteobacterial ancestor, *Rickettsia prowazekii* [1,2] that were harnessed by a eukaryotic cell as the host billions of years ago [3]. The proteobacterium became a symbiote of the host cell, bringing with it a system for more efficient generation of cellular energy in the form of ATP.

During the course of evolution, the genetic material of the mitochondria underwent reductive expansion and was transferred in a retrograde fashion to the nuclear genome. The retrograde movement of genes from the mitochondria to its eukaryotic host paved the way for the mitochondria to specialize as energy production organelles rather than consuming energy repetitively replicating its own multi-copy genome. The mitochondrial-encoded genetic material at present is a vestige of the original proteobacterial genome [4–6] meaning despite having DNA of their own, mitochondria rely heavily on the nuclear genes for most of their functions.

Mitochondria have critical functions in metabolism, organ homeostasis, apoptosis and aging. They also play important but still largely mysterious roles in human pathology, as demonstrated by the enormous biological variation and diverse disorders in patients with mitochondrial disease [7–10]. Nearly every organ system can be compromised, but with highly variable and complex physiological and biochemical outcomes. Imaging and basic science of mitochondria showcase how this highly dynamic organelle responds differentially to extrinsic and intrinsic biological signals. However, understanding how mitochondria function in normal biology, and how human mitochondrial DNA variations contribute to health and disease, has been hampered by a lack of effective approaches to manipulate the powerhouse of the cell.

The vertebrate mitochondrial chromosome is circular and includes 37 genes, 13 encoding for protein subunits of the electron transport chain, 22 coding for transfer RNAs, and 2 encoding ribosomal RNAs **(Figure 1)** [11,12]. The mitochondrial gene order, strand specific nucleotide bias and codon usage is highly conserved [13]. However, mtDNA encoded genes lack introns and utilize a divergent genetic code than their nuclear counterparts [14,15]. For instance, AUA codon codes for methionine as per mitochondrial genetic code, whereas the same sequence codes for isoleucine in the nuclear genetic code. Similarly, nuclear stop codon UGA is read as the tryptophan amino acid by the mitochondrial codon cypher.

**Figure 1:**
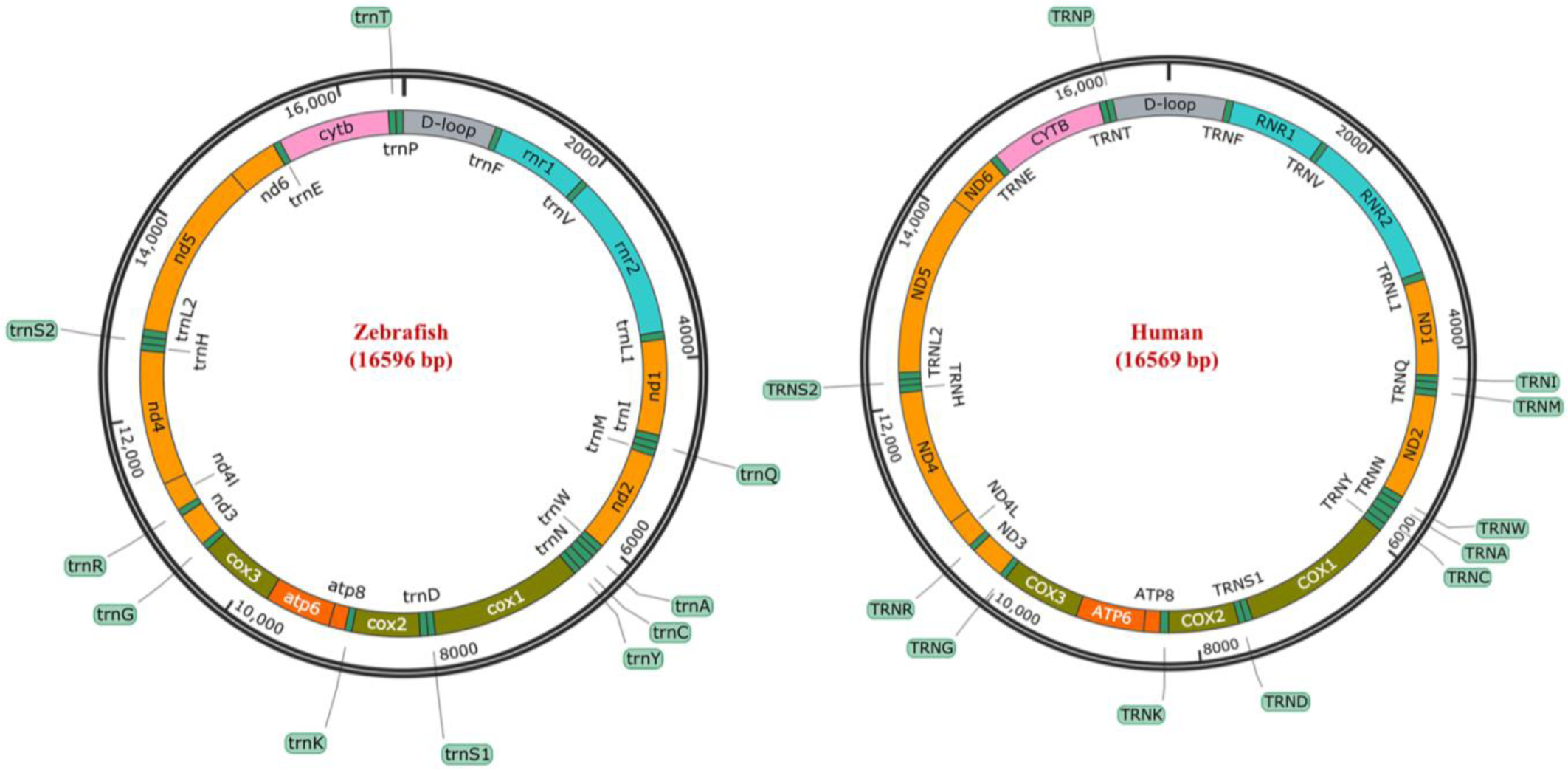
Circular representation of the zebrafish (NC_002333.2) and human mitochondrial genomes (NC_012920.1): Both genomes share the same synteny, number of genes and are nearly identical in size.

Mitochondria are unique cellular compartments with different DNA and RNA repair and editing rules, hampering attempts at directly manipulating these nucleic acid components. For example, DNA nucleases that introduce double stranded breaks and subsequent repair in nuclear DNA induce the degradation of mtDNA [16–18]. Indeed, none of the canonical DNA repair pathways found in the nucleus have been shown to be active in mitochondria [18–20]. Finally, no system has demonstrated the ability to deliver exogenous DNA or RNA to mitochondria, restricting the tools available for mtDNA editing [21,22]. All of these factors are distinct from the nuclear genome, making mtDNA a far less accessible genome for traditional gene editing methods and reagents.

The diverse functions that mitochondria are capable of, including oxidative phosphorylation, would not be possible with the small subset of 13 proteins encoded in mtDNA. Discoveries led by high throughput proteomic approaches have enhanced our knowledge of the mitochondrial proteome [23–27]. There are approximately 1158 nuclear-encoded proteins that localize to the mammalian mitochondria, exerting a dual genetic control via its nuclear counterpart. Nuclear-encoded mitochondrial proteins on the basis of their mitochondrial function can be broadly classified into different categories including oxidative phosphorylation, energy production, membrane dynamics, genome maintenance, and ion/metabolite homeostasis.

Unlike the nuclear genome, where most cells have only two copies, each cell can harbor thousands of copies of mtDNA, depending on environmental needs, making mitochondrial genome engineering a population genetics challenge. When every mtDNA molecule is identical with non-mutagenic variants, the cell is in a genetic state, called homoplasmy, and the mitochondria functions normally. In people with mtDNA-based mitochondrial disease, mtDNA genomes harboring pathogenic mutations typically co-exist with the healthy genomes in a state called heteroplasmy. Homoplasmy may also arise when the cell harbors only mutant mtDNA molecules, resulting in severe clinical manifestations. [28]. The ratio of pathogenic-harboring to nonpathogenic-harboring mtDNA is critical to mitochondrial disease onset. A threshold exists for most mitochondrial disease, at which, the healthy mtDNA genomes can no longer compensate for the pathogenic genomes, resulting in disease onset leading to breakdown of oxidative phosphorylation and other mitochondrial function(s) [29].

Mitochondrial disorders are a heterogeneous group of clinical manifestations resulting from either inherited or spontaneous mutations in mtDNA or nDNA leading to altered structure or function of the proteins or RNA that reside in mitochondria [6,10,30,31]. The pathophysiology of mitochondrial disorders involves functions of both the nuclear and mitochondrial genomes, conferring additional complexity to the manifestation of the syndromes. The prevalence of mitochondrial disorders is about 1 in 5000 live births; however, the prevalence is often influenced by the presence of founder mutations and consanguinity in populations [31–33]. These conditions range in severity dependent on the specific mutation and its prevalence and localization within the patient. In addition to diseases specifically progenerated by mitochondrial defects, mitochondrial dysfunction has also been implicated as a cofactor or by-product of many other conditions; including cancer, immune diseases, developmental delays and neurodegenerative condition such as Alzheimer’s [34–38].

Mitochondrial dysfunction tends to affect primarily high energy systems and can therefore have devastating effects on a wide range of body systems including brain function, liver function, vision, hearing, immune function, and all muscle types [31,39]. The identification and characterization of novel mitochondrial genes in human genetic disorders have also enhanced our knowledge of mitochondrial function [40–44]. However, despite their known prevalence and severity, mitochondrial diseases remain understudied as compared to other genetic conditions, and the options for treatment are limited.

The conserved roles of mitochondria have been traditionally uncovered using accessible model systems such as the yeast *S.cerevisiae* [45–48] and invertebrates including *C.elegans* [49–52] and *D. melanogaster* [53–56]. Vertebrate mitochondria, however, are known to have additional innovations and encode unique subunits not present in invertebrate models [5,45,57–67]. Most exploration to date of vertebrate mitochondrial biology has been conducted either in cell culture or in mouse models. However, cells in a dish are missing important environmental cellular and organismal context, whereas mouse mitochondrial experimental work can be hindered by limited population size and invasive imaging techniques.

We provide here an initial genetic toolkit for modeling mitochondrial biology and disease in *Danio rerio* (zebrafish) in an effort to help catalyze the use of this invaluable model organism in mitochondrial research. The zebrafish is a tropical freshwater teleost that has proven to be an invaluable resource for the study of human disease and genetics [68–70]. Principal among the many research amenable characteristics of zebrafish are their high genetic orthology to humans, high fecundity, their optical transparency (in the embryonic stage) that lends itself to facile imaging, and the ease of reagent delivery through microinjection into the single celled embryo, which is a full millimeter across before the first mitotic division. Zebrafish also serve as a potentially powerful vertebrate model organism to study human mitochondrial disorders because of conserved mitochondrial genome and mitochondrial genetic machinery. Zebrafish and human mitochondrial chromosomes display ∼65% sequence identity at the nucleotide level, and share the same codon usage, strand specific nucleotide bias and gene order [12] **(Figure 1)**.

Over the recent years, zebrafish research has helped shed light on mitochondrial biology and further developed our understanding of the mechanisms of mitochondrial-associated pathology [71–76]. Zebrafish have also been successfully used as a model system to study mitochondrial targeting drugs with implications in development and cardiovascular function [71,77,78]. These drugs can simply be dissolved in the water housing the larvae and offer an advantage of large sample size with minimal volumes of drug administered. Drug-screening studies have aided in understanding the pathogenesis of various diseases and helped to identify targets for treatment [77].

To help engender an expansion of zebrafish deployment in mitochondrial research, we present a panel or “starter-pack” toolkit of genetic mutants made in nuclear-encoded mitochondrial genes in zebrafish. This mutant panel, which we have named the Marriot Mitochondrial Collection (MMC), consists of 28 zebrafish lines with mutations in 23 different mitochondrial genes. The mutant collection focusses primarily on genes that encode for components of the energy generating electron transport chain with at least one mutant in each complex of the ETC. Other mutants consist of assembly factors, protein chaperones that manage mitochondrial membrane traffic, and genes related to mitochondrial replication. These mutants were made either by the use of targeted endonuclease [79], or curated from our research group’s library of randomly generated insertional mutants [80]. We hope that this collection, in addition to being intrinsically useful, will also help serve as a primer to the modeling of mitochondrial biology and disease in zebrafish.

## Methods

### Zebrafish Handling

All animal work was conducted under Mayo Clinic’s institutional animal welfare approvals (IACUC number: A34513-13-R16).

### Identification of zebrafish orthologs having putative mitochondrial function

A previous study combining discovery and subtractive proteomics with computational, microscopy identified 1098 mouse genes that could encode for proteins residing in mitochondria [24]. They further identified 1013 human orthologs for these genes, providing an initial inventory of the genes coding for proteins resident in mitochondria. Using literature assessment and HUGO database curation approaches, we identified 97 proteins that are involved in the biogenesis and assembly of electron transport chain in mitochondria. Using zebrafish orthologs of human genes from ZFIN (Zebrafish Information Network), 93 zebrafish mitochondrial orthologs were identified. These orthologs were systemically annotated with respect to clinical phenotype by extensive mining from PubMed based published case reports and OMIM database **(Supplementary Table 1)**.

### Mutant Generation

Mutant lines were created by one of two methods, either through the targeted use of Transcription Activator Like Effector Nucleases (TALENs) [79] or by screening Gene Breaking Transposon (GBT) lines for integrations into mitochondrial genes [80]. Reagents were delivered in both methods by the microinjection of either TALEN pairs or GBT transposon and Tol2 transposase into single cell zebrafish embryos.

### TALEN Design/Assembly/Delivery

The TALEN mutants in this collection were originally generated as part of a previously published study [79]. In brief, TALEN pairs were designed using the Mojohand software platform [81] (www.talendesign.org) to target highly conserved, and therefore likely functionally important, areas of nuclear-encoded mitochondrial genes. TALEN RVDs were then cloned into pT3Ts-GoldyTALEN (TALEN vector with a T3 transcriptional promoter for in vitro transcription) using the FusX rapid TALEN assembly system [79], which uses the RVD definitions: HD=C, NN=G, NI=A, NG=T. Following assembly, mRNA was synthesized in vitro using the mMessage Machine T3 kit (Ambion) and extracted by a phenol-chloroform extraction as prescribed in the mMessage Machine manual. The extracted mRNA was then delivered into single cell zebrafish embryos at 100pg doses (50pg per TALEN arm) by microinjection.

### TALEN Mutant Screening

Following microinjection, genomic DNA was extracted from F0 larvae three days post fertilization (dpf) by sodium hydroxide extraction. DNA for eight individual larvae was analyzed for NHEJ activity at the TALEN target site by Restriction Fragment Length Polymorphism (RFLP) analysis. Groups with high reported NHEJ activity by RFLP were raised to adulthood and outcrossed to create an F1 generation. Suspected NHEJ mutants were further verified by Sanger sequencing. F1 larvae demonstrating NHEJ mutations confirmed by both RFLP and Sanger sequencing were raised to adulthood. Fin biopsies were performed on these adults and DNA was extracted by sodium hydroxide extraction.

### Gene Breaking Transposon System

The Gene Breaking Transposon System of protein trap system and a complete repository of protocols for the creation and screening of GBT mutant lines has been described [80]. In short, protein trap transposons were delivered in combination with mRNA for Tol2 transposase (25pg each) into single cell zebrafish embryos by microinjection. Embryos were screened for GFP fluorescence at 3-4dpf and classified broadly into three classes [80]. Class three embryos, those with whole body GFP expression, were raised to adulthood and outcrossed to non-transgenic lines to create an F1 generation. mRFP-expressing F1 embryos were sorted by expression pattern, assigned a GBT number, and raised to adulthood. These adult fish were then outcrossed to non-transgenic lines to create an F2 generation upon which all subsequent propagation, testing and imaging was conducted.

To determine genes tagged by the protein trap system, rapid <u>a</u>mplification of <u>c</u>DNA <u>e</u>nds (RACE) was performed as described [82] with minor updates to primer sequences. cDNA was generated using a transposon-specific primer (5R-mRFP-P0) against 250ng of total mRNA in the reverse transcription reaction. PCR was then performed using the following gene-specific primers: 5R-mRFP-P1 and 5R-mRFP-P2. The resulting products were TA cloned for further amplification and then sequence verified for in-frame mRFP fusions by Sanger sequencing.

In some cases, inverse PCR was also conducted to RACE PCR as described [80]. Whole genomic DNA was extracted from individual F2 embryos using a sodium hydroxide extraction and 800ng was digested in a combination reaction using AvrII, NheI, SpeI, and XbaI restriction enzymes. Approximately 200ng of the product of this digestion was self-ligated and used as a template for PCR using the following nested and primary primers. 5’ side: 5R-mRFP-P1 and 5R-mRFP-P2 paired with INV-OPT-P1 and INV-OPT-P2, respectively. 3’ side: 5R-GFP-P1 and 5R-GFP-P2 with Tol2-ITR(L)-O1 and Tol2-ITR(L)-O3, respectively. The products of the final nested reaction were gel-extracted, cloned, and sequenced. Prospective in frame mRFP fusions were further verified by comparing suspected protein trap allele to GBT expression pattern and by PCR against DNA or cDNA from mRFP carrier siblings versus non-carrier siblings.

Protein trap lines as verified by these methods were catalogued for specific genomic fusions using the National Center for Biotechnology Information’s (NCBI) Homologene Database. Human orthologs for tagged genes were further identified using blastX searches against the human genome. GBT lines with protein trap fusions to nuclear encoded mitochondrial genes with human orthologues have been included in this collection.

### *In silico* analysis to determine protein homology and alteration of DNA sequence by TALENs and GBT in zebrafish mutants

Human amino acid sequences were compared to both the zebrafish wild type sequence and each specific zebrafish mutant sequence associated with each allele. Amino acid sequence information for a particular gene was gathered for humans from https://www.uniprot.org/ along with the wild type amino acid sequence for the zebrafish. The first/most common isoform for each entry was used for this analysis. Any added information regarding functional domains or regions was also gathered. A protein-protein BLAST was conducted using https://blast.ncbi.nlm.nih.gov/Blast.cgi to compare the sequences of the human and wild type zebrafish amino acid sequences. BLAST search settings used included the blastp algorithm. Regions of low and high homology were then mapped out for the sequences.

For zebrafish mutant to human comparisons, the cDNA or DNA sequences were gathered via sequencing and put through the translate tool at https://web.expasy.org/translate/. The standard genetic code was used for conversion. Stop codons arising from the frameshifted DNA sequence were found in all mutants. The BLAST analysis conducted on the mutants was taken from the starting methionine to the first encountered stop codon. Regions of low and high homology were then mapped for each sequence.

### Access to all reported reagents – zebrafish and sequences

All listed tools are immediately available through the Mayo Clinic Zebrafish Facility, and all fish lines will be available via ZIRC. Sequences needed for genotyping and related metadata are currently on zfishbook [83].

## RESULTS

### Generation of zebrafish mutant collection

We generated zebrafish mutants for a wide selection of nuclear-encoded mitochondrial genes by employing two genetic engineering approaches, gene breaking transposons and TALENs. The collection of zebrafish mutants is referred as the Marriot Mitochondrial Collection (MMC) and comprises of 28 mutants for 23 nuclear-encoded mitochondrial proteins **(Figure 2)**. Out of these, 22 were created by TALEN indel mutagenesis and 6 were created by gene breaking trap mutagenesis. The mutants include proteins from all known functional pathways involved in mitochondrial homeostasis and ATP generation **(Figure 2)**.

**Figure 2:**
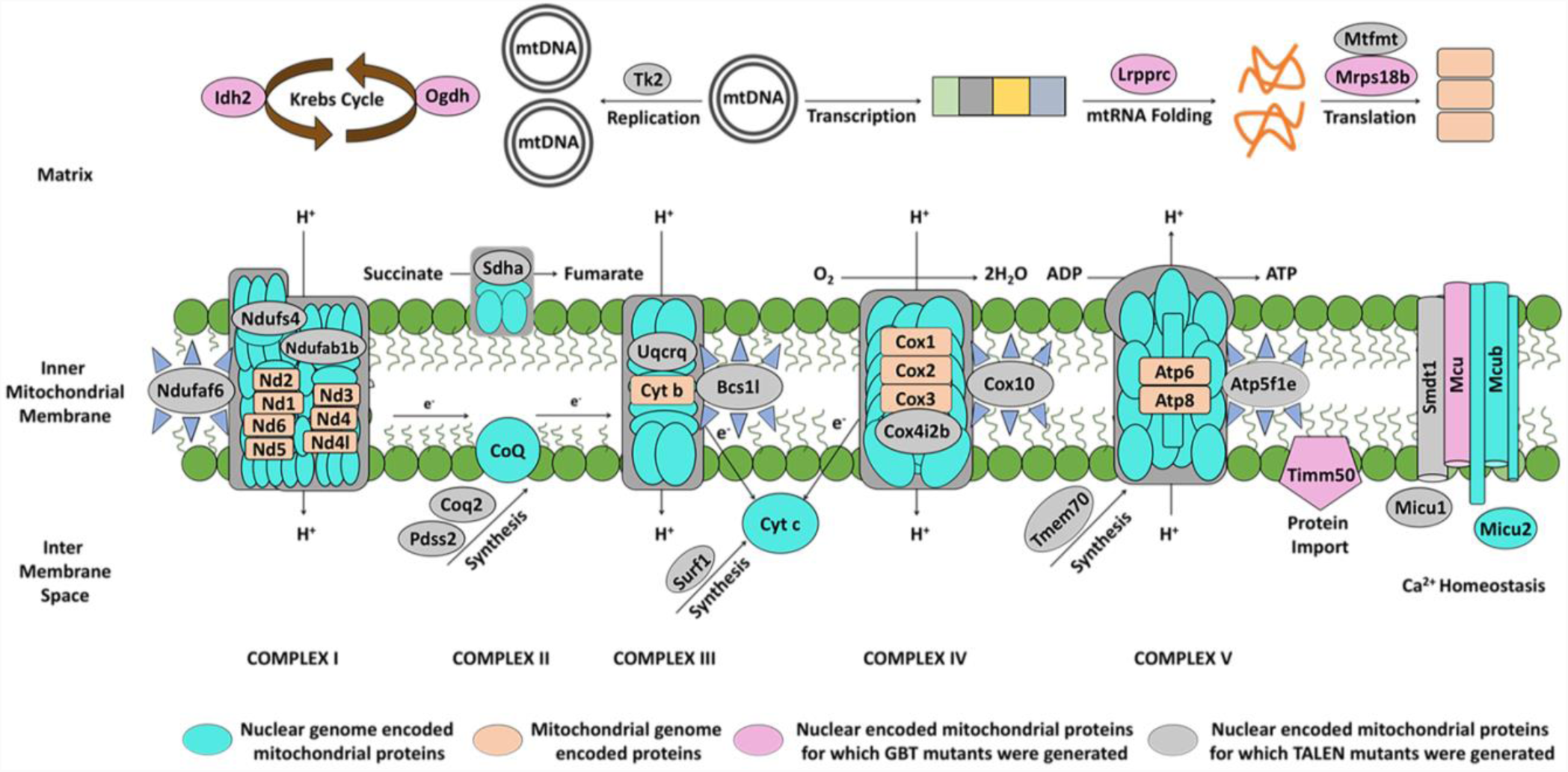
Zebrafish Marriot mitochondrial mutant collection: Schematic representation of various mitochondrial resident pathways for which zebrafish mutants were generated. The nuclear-encoded mitochondrial proteins have been illustrated according to the function they are involved in mitochondrial maintenance and homeostasis. Mutants generated by gene editing are depicted as grey, whereas those generated by the GBT system are depicted in pink.

Broadly, the pathways can be classified as subunits of oxidative phosphorylation complexes, chaperones for assembly of oxidative phosphorylation proteins, maintenance proteins for the mitochondrial genome (replication, mtRNA folding and translation), calcium homeostasis and mitochondrial protein import. All of these genes have human orthologs, nearly all with known mutations in which lead to severe clinical manifestations such as Leigh syndrome, cardiomyopathy, progressive external ophthalmoplegia, and oxidative phosphorylation deficiency **(Table 1)**.

**Table 1:** List of nuclear-encoded genes for which zebrafish mutants were generated as part of MMC collection: The current MMC resource list summarizes information on human gene, zebrafish ortholog, mouse ortholog, relevant clinical phenotypes and diseases, protein function, and OMIM ID. Green = Created by TALEN Indel Mutagenesis, Orange = Curated from GBT protein trap lines. Allele designators are highlighted in bold for respective zebrafish paralog. (Chr-Chromosome; OMIM ID: Online Mendelian Inheritance in Man ID; NA-not available).

| Approved symbol | Approved name | Human chr | Zebrafish orthologs | Zebrafish chr | Allele designator | Mouse orthologs | Mouse chr | Clinical phenotype observed in humans | Disease | Biological function | OMIM ID |
| --- | --- | --- | --- | --- | --- | --- | --- | --- | --- | --- | --- |
| *NDUFAB1* | NADH:ubiquinone oxidoreductase subunit AB1 | 16 | *ndufab1a*<br>***\*ndufab1b*** | 1<br>3 | **\*mn0135** | Ndufab1b | 7 | -na- | -na- | Oxidative phosphorylation enzymes - Complex I | 603836 |
| *NDUFA6* | NADH:ubiquinone oxidoreductase complex assembly factor 6 | 8 | *ndufaf6* | 16 | mn0117 | Ndufaf6 | 4 | Focal right-hand seizures, ataxia, lactic acidosis, exercise intolerance, weakness and muscle tension | Leigh syndrome, mitochondrial complex I deficiency | Oxidative phosphorylation complex assembly - Complex I | 612392 |
| *NDUFS4* | NADH:ubiquinone oxidoreductase subunit S4 | 5 | *ndufs4* | 5 | mn0118 | Ndufs4 | 13 | Mitochondrial complex I deficiency | Mitochondrial complex-I deficiency | Oxidative phosphorylation enzymes - Complex I | 602694 |
| *SDHA* | Succinate dehydrogenase complex flavoprotein subunit A | 5 | *sdha* | 19 | mn0121 | Sdha | 13 | Dyspnea, cardiomegaly, cardiomyopathy, nystagmus, hypotonia, gastrointestinal stromal tumors, paragangliomas, pheochromocytoma, psychomotor regression and severe hyperandrogenism | Mitochondrial complex-II deficiency; Cardiomyopathy, Leigh syndrome; Paraganglioma | Oxidative phosphorylation enzymes - Complex II | 600857 |
| *BCS1L* | BCS1 homolog, ubiquinol-cytochrome C reductase complex chaperone | 2 | *bcs1l* | 9 | mn0102 | Bcs1l | 1 | Growth retardation, aminoaciduria, cholestasis, iron overload, lactic acidosis, early death, hypotonia, encephalopathy, microcephaly, psychomotor regression, hypoglycemia and hepatic failure | Björnstad syndrome; GRACILE syndrome; Leigh syndrome; Mitochondrial complex III deficiency, nuclear type 1 | Oxidative phosphorylation complex assembly - Complex III | 603647 |
| *UQCRCQ* | ubiquinol-cytochrome c reductase complex III subunit VII | 5 | *uqcrcq* | 14 | mn0128<br>mn0129 | Uqcrcq | 11 | Severe psychomotor retardation and extrapyramidal signs, dystonia, athetosis, ataxia, mild axial hypotonia and marked global dementia | Mitochondrial complex III deficiency | Mitochondrial complex III: ubiquinol-cytochrome c reductase complex subunits | 612080 |
| *COX10* | Cytochrome c oxidase assembly factor heme A: farnesyltransferase COX10 | 17 | *cox10* | 12 | mn0107<br>mn0108 | Cox10 | 11 | Muscle weakness, hypotonia, ataxia, ptosis, pyramidal syndrome, status epilepticus, lactic acidosis, hypertrophic cardiomyopathy, hypoglycemia and metabolic acidosis | Leigh syndrome due to mitochondrial COX IV deficiency; Mitochondrial complex IV deficiency | Oxidative phosphorylation complex assembly - Complex IV | 602125 |
| *COX4I2* | Cytochrome C Oxidase Subunit 4I2 | 20 | *cox4i2b* | 23 | mn0104 | Cox4i2 | 2 | Exocrine pancreatic insufficiency, dyserythropoietic anemia, and calvarial hyperostosis | Exocrine pancreatic insufficiency, dyserythropoietic anemia, and calvarial hyperostosis | Oxidative phosphorylation complex enzymes - Complex IV | 607976 |
| *SURF1* | SURF1, Cytochrome C Oxidase Assembly Factor | 9 | *surf1* | 5 | mn0123 | Surf1 | 2 | Childhood onset neuropathy, lactic acidosis, mild intellectual disability ataxia, facial dysmorphism, encephalopathy, hypotonia, cerebellar ataxia, deafness, ophthalmoplegia, growth retardation and nystagmus | Charcot-Marie-Tooth disease, type 4K; Leigh syndrome, due to COX IV deficiency | Involved in biogenesis of cytochrome c oxidase complex | 185620 |
| *ATP5F1E* | ATP synthase F1 subunit epsilon | 20 | *atp5f1e* | 6 | mn0101 | Atp5f1e | 2 | Neonatal-onset lactic acidosis, 3-methylglutaconic aciduria, mild mental retardation and peripheral neuropathy | Mitochondrial complex (ATP synthase) deficiency | Oxidative phosphorylation complex assembly - ATP synthase | 606153 |
| *TMEM70* | Transmembrane protein 70 | 8 | *tmem70* | 2 | mn0126<br>mn0127 | Tmem70 | 1 | Lactic acidosis, encephalopathy, histiocytoid cardiomyopathy, microcephaly, hypotonia, facial dysmorphism and 3-methylglutaconic aciduria psychomotor delay and hyperammonemia | Mitochondrial complex V (ATP synthase) deficiency | Biogenesis of mitochondrial ATP synthase | 612418 |
| *COQ2* | Coenzyme Q2, Polyprenyltransferase | 4 | *coq2* | 5 | mn0106 | Coq2 | 5 | Multiple system atrophy, unsteadiness of gait, nystagmus, gait ataxia, dysarthria, speech difficulty, dysmetria, lactic acidosis, urinary dysfunction and nystagmus | Coenzyme Q10 deficiency, primary; Multiple system atrophy | Biosynthesis of CoQ, coenzyme in mitochondrial respiratory chain | 609825 |
| *PDSS2* | Decaprenyl diphosphate synthase subunit 2 | 6 | *pdss2* | 13 | mn0120 | Pdss2 | 10 | Hypotonia, seizures, cortical blindness, lactic acidosis, encephalopathy and nephrotic syndrome | Coenzyme Q10 deficiency | Involved in biosynthesis of coenzyme Q | 610564 |
| *MICU1* | Mitochondrial calcium uptake 1 | 10 | *micu1* | 13 | mn0132 | Micu1 | 10 | Proximal muscle weakness and learning disabilities | Myopathy with extrapyramidal signs | Regulator of mitochondrial calcium uptake | 605084 |
| *SMDT1* | Single-pass membrane protein with aspartate rich tail | 22 | *smdt1a*<br>*\*smdt1b* | 3<br>1 | *\*mn0122* | Smdt1 | 15 | -na- | -na- | Core regulatory subunit of calcium channel in mitochondrial inner membrane | 615588 |
| *MCU* | Mitochondrial calcium uniporter | 10 | *mcu* | 13 | mn0111 | Mcu | 10 | -na- | -na- | Calcium transporter and mediates calcium uptake in mitochondria | 614197 |
| *TK2* | Thymidine kinase 2, mitochondrial | 16 | *tk2* | 7 | mn0125 | Tk2 | 8 | Infantile onset fetal encephalomyopathy, failure to thrive, muscle weakness, mild facial weakness, myopathy, axonal neuropathy, brain atrophy, hypertrophic cardiomyopathy, regression of motor development, psychomotor arrest, dysphonia, dysphagia, ptosis, impaired gait, bilateral ptosis, hypotonia, lactic acidosis and epilepsy | Progressive external ophthalmoplegia; Mitochondrial DNA depletion syndrome 2 (myopathic type) | Required for mitochondrial DNA synthesis | 188250 |
| *MTFMT* | Mitochondrial methionyl-tRNA formyltransferase | 15 | *mtfmt* | 7 | mn0113<br>mn0114 | Mtfmt | 9 | Psychomotor developmental delay, renal dysplasia, mild facial dysarthria and ataxia | Combined oxidative phosphorylation deficiency | Catalyzes the formylation of methionyl-tRNA | 611766 |
| *MCU* | Mitochondrial calcium uniporter | 10 | *mcu* | 13 | mn0599gt | Mcu | 10 | -na- | -na- | Calcium transporter and mediates calcium uptake in mitochondria | 614197 |
| *LRPPRC* | Leucine rich pentatricopeptide repeat containing | 2 | *lrpprc* | 13 | mn0235gt | Lrpprc | 17 | Delayed psychomotor development, mental retardation, mild dysmorphic facial features, hypotonia, ataxia, development of lesions in the brainstem and basal ganglia, seizures, dysphagia and hypertrophic cardiomyopathy | Leigh syndrome, French-Canadian type | Involved in translation of mitochondrial encoded cox subunits and mediation of folding of mitochondrial transcriptome | 607544 |
| *MRPS18B* | Mitochondrial ribosomal protein S18B | 6 | *mrps18b* | 19 | mn0425gt | Mrps18b | 17 | -na- | -na- | Part of small 28S subunit of mitoribosome | 611982 |
| *TIMM50* | Translocase of inner mitochondrial membrane 50 | 19 | *timm50* | 15 | mn0906gt | Timm50 | 7 | Severe intellectual disability, seizure and 3-methylglutaconic aciduria | 3-methylglutaconic aciduria, type IX; Mitochondrial complex V deficiency | Subunit of TIM23 inner mitochondrial membrane complex and recognizes mitochondrial targeting signal or pre-sequence | 607381 |
| *OGDH* | Oxoglutarate dehydrogenase | 7 | *ogdha*<br>*\*ogdhb* | 8<br>10 | *mn0281gt | Ogdh | 11 | Colorectal cancer | Alpha-ketoglutarate dehydrogenase deficiency | Catalyzes the conversion of 2-oxoglutarate to succinyl-CoA and CO<sub>2</sub> | 613022 |
| *IDH2* | Isocitrate dehydrogenase (NADP (+)) 2, mitochondrial | 15 | *idh2* | 18 | mn0268gt | Idh2 | 7 | Acute myeloid leukemia and abnormal production of D-2-hydroxyglutaric acid | D-2-hydroxyglutaric aciduria 2 | Catalyzes the conversion of isocitrate to 2-oxoglutarate | 147650 |

### TALEN- and GBT-mediated targeting of nuclear-encoded mitochondrial genes

Amino acid analysis using protein-protein BLAST functions between the wild type human, wild type zebrafish, and mutant zebrafish sequences showed consistent results between the three conditions. The comparison of wild type human and wild type zebrafish sequences showed high areas of homology following the mitochondrial targeting domain in almost every gene, with 80-100% similarity in catalytic or active domains of the transcripts. Analysis of the mutant zebrafish and human wild type comparison showed a range of difference with predicted frameshift mutations leading to truncation of the protein very early in the transcript **(Figure 3A-X)**. Of the alleles created by TALEN mutagenesis in the MMC collection, all but one (*micu1)* showed a predicted frameshift mutation leading to a truncation event immediately following the NHEJ-mediated insertion or deletion. The GBT mutants showed high levels of homology prior to the transposon integration site followed by a nearly complete loss of normal transcript levels following splicing into the GBT cassette.

**Figure 3:**
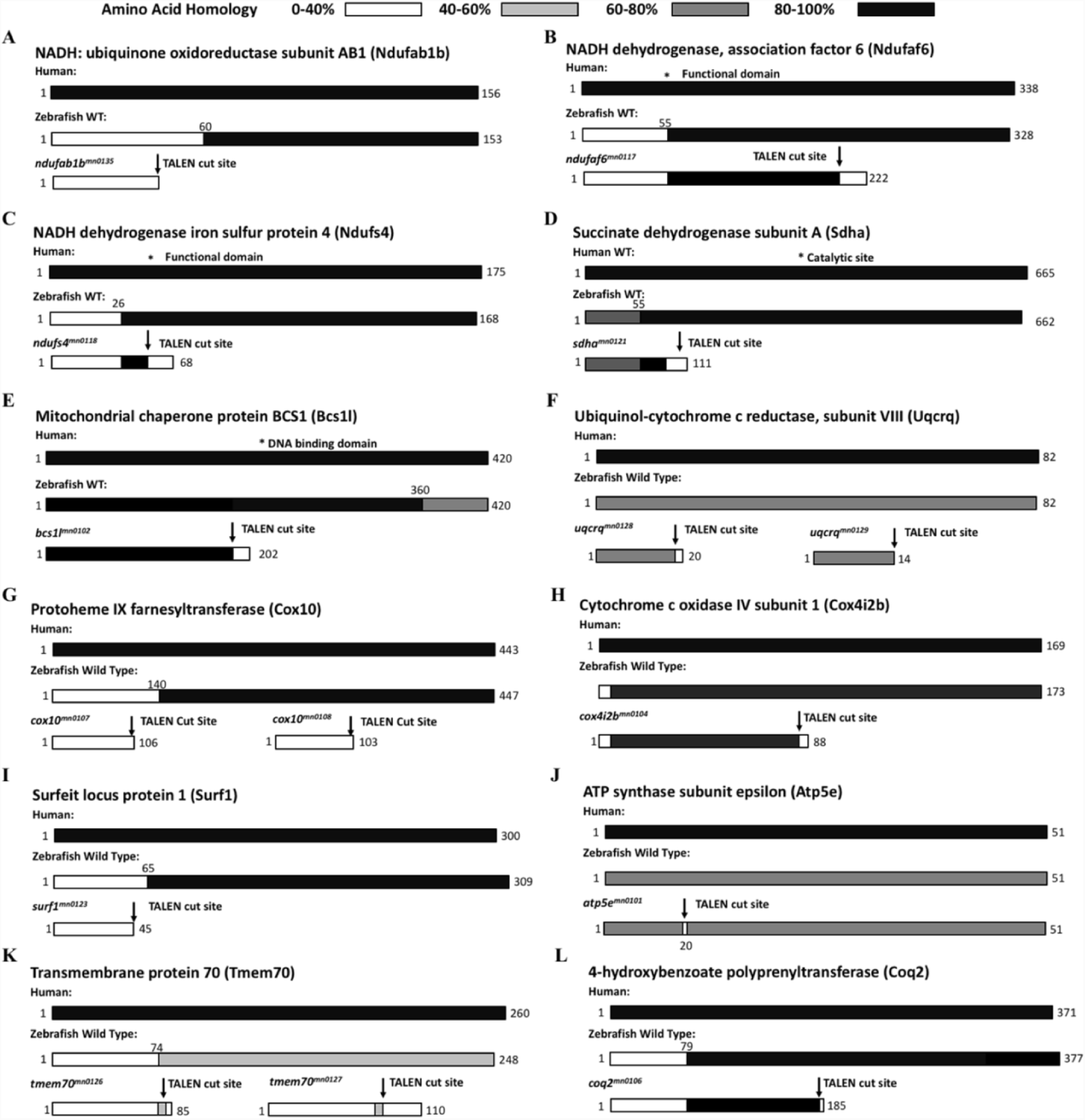
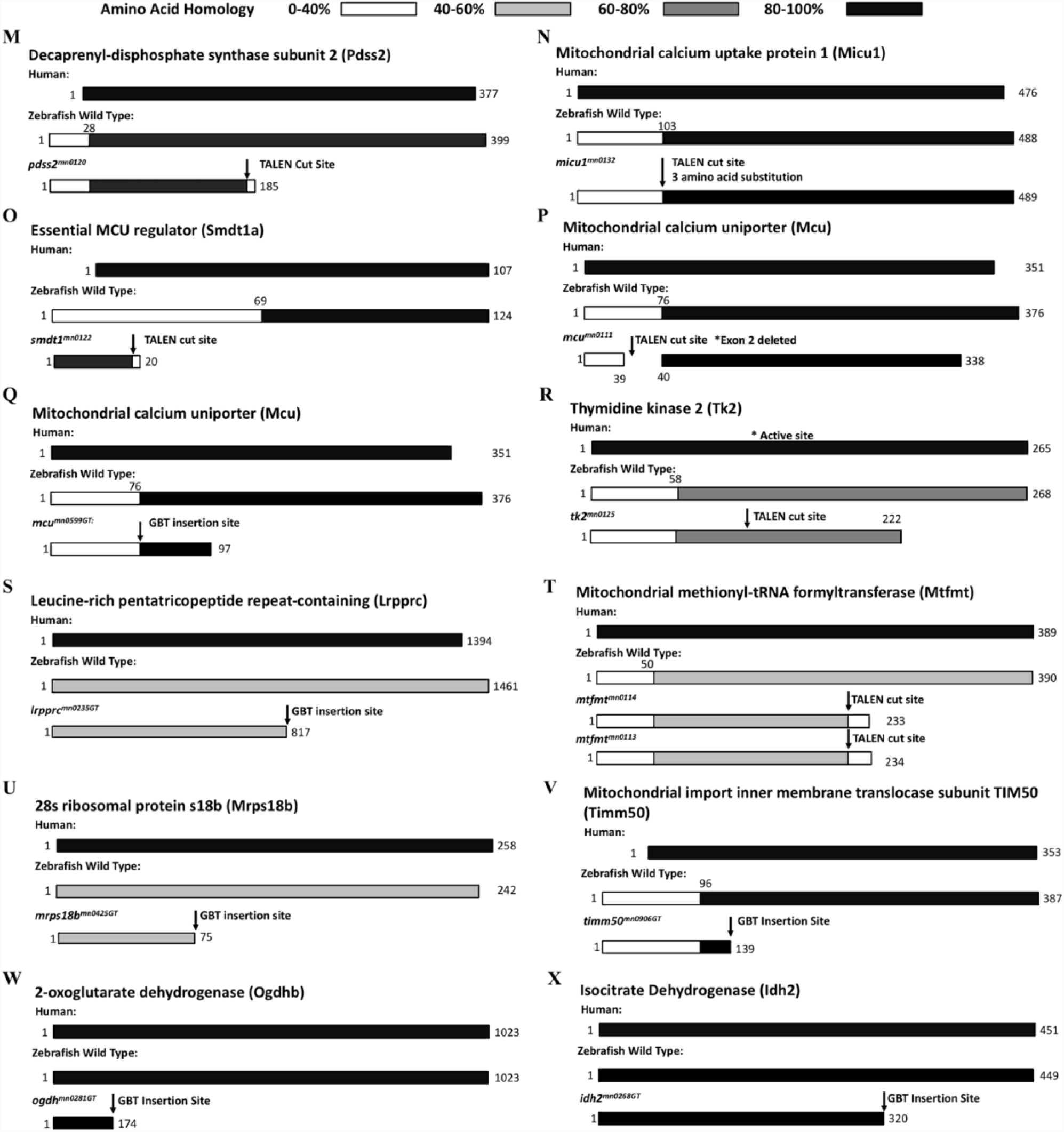
A depiction of homology of the mutants created for proteins involved in different mitochondrial resident pathways: A-C: Complex 1 of mitochondrial respiratory chain; A: NADH: ubiquinone oxidoreductase subunit AB1 b (Ndufab1b); B: NADH dehydrogenase association factor 6 (Ndufaf6); C: NADH dehydrogenase iron sulfur protein 4 (Ndufs4); **D: Complex 2 of mitochondrial respiratory chain,** Succinate dehydrogenase subunit A (Sdha), subunit; **E-F: Complex 3 of mitochondrial respiratory chain;** E: Mitochondrial chaperone protein BCS1 (Bcs1l); F: Ubiquinol-cytochrome c reductase subunit VIII (Uqcrq); **G-I: Complex 4 of mitochondrial respiratory chain;** G: Protoheme IX farnesyltransferase (Cox10); H: Cytochrome c oxidase IV subunit 1 (Cox4i2b); I: Surfeit locus protein 1 (Surf1); **J-K: Complex 5 of mitochondrial respiratory chain;** J: ATP synthase subunit epsilon (Atp5e); K: Transmembrane protein 70 (Tmem70); **L-M: Biosynthesis of coenzyme Q;** L: 4-hydroxybenzoate polyprenyltransferase (Coq2) M: Decaprenyl-disphosphate synthase subunit 2 (Pdss2); **N-Q: Mitochondrial calcium homeostasis;** N: Mitochondrial calcium uptake protein 1 (Micu1); O: Calcium uniporter protein (Mcu) generated by TALEN and P; GBT; Q: Essential MCU regulator a (Smdt1a); **R-U: Mitochondrial genome maintenance;** R: Thymidine kinase 2 (Tk2); S: Leucine-rich pentatricopeptide repeat-containing (Lrpprc); T: Mitochondrial methionyl-tRNA formyltransferase (Mtfmt); U: 28s Ribosomal protein s18b, mitochondrial (Mrps18b); **V: Mitochondrial protein import;** Mitochondrial import inner membrane translocase subunit TIM50 (Timm50); **W-X: Mitochondrial metabolite synthesis;** W: 2-Oxoglutarate dehydrogenase b (Ogdhb); X: Isocitrate dehydrogenase (Idh2).

## Discussion

In a multicellular organism, each cell is able to carry out its functions due to well-orchestrated cross-talk between the nuclear and mitochondrial genomes. Many mitochondrial functions like energy production, genome maintenance, ion/metabolite homeostasis, membrane dynamics and transport of biomolecules can be attributed to approximately 1158 known nuclear proteins residing in the mitochondria [24,27]. Due to advancements in proteomic technologies in recent years, there has been a surge in documentation of many characterized and uncharacterized mitochondrial proteins. However, correlation between mitochondrial localization of these proteins and their physiological significance in disease progression remains largely unexplored. Out of the 1158 nuclear encoded proteins, only 245 have functional evidence in mitochondrial clinical manifestations [39]. For the remaining 913 proteins, mitochondrial involvement in disease progression has yet to be demonstrated. The obstacles in mitochondrial genetic research, and thus delays in finding effective treatments, are primarily due to the limited tools available to mitochondrial researchers, specifically the small number of available model systems and animals.

Most mitochondrial research thus far has made use of model systems that each present their own unique challenge to accurate study of human disease. Yeast, one of the most common laboratory eukaryotes, have been incredibly useful in mitochondrial research, but unlike humans and other animals they lack complex I [45]. Complex 1 deficiency is the most frequent among mitochondrial disorders caused by mutations in 28 out of 48 genes contributing to its assembly and biogenesis [84]. Also, complex 1 is responsible for the most ROS generation, which acts as a cue in various signaling pathways [85].

Human cell culture has been valuable in exploring mitochondrial disease models, principally through cytoplasmic hybrids that are made by replacing the mitochondria of an immortalized cell line with mutated patient mitochondria [86–89]. However, while these cells are a more faithful representation of human mitochondrial activity than yeast, they have different barriers to accuracy. First, immortalized cells, and cultured cells in general, often have modified metabolic pathways compared to cells *in vivo*. Second, as is common to all cell culture research, cells in a dish cannot accurately represent the complex interactions and systems biology inherent to a complete organism.

In this study, we aim to support the establishment of the zebrafish as a complementary animal model for understanding the role of nuclear-encoded mitochondrial proteins in biology and the pathophysiology of mitochondrial disorders. We propose that zebrafish can be an excellent model organism to study mitochondrial biology primarily because of their conserved genome, codon bias and synteny. Other advantages include their amenability to genetic manipulation and optical clarity, which facilitates direct observation. New gene editing techniques such as TALENs and CRISPRs have aided in the development of humanized disease models in organisms such as zebrafish, mice, and even pigs [90–97].

The MMC collection described here encompasses mutations in 23 different nuclear-encoded genes related to mitochondrial function enabling a diverse study of the important roles the mitochondria play in cellular biology. The collection includes mutants in all five complexes of the electron transport chain as well as many other crucial pathways related to protein transport, metabolite synthesis, mtDNA replication and expression, and calcium homeostasis. The first set of mutants were created using custom gene editing to exons coding for critical functional domains of the protein product. These domains were targeted because they shared high levels of homology to their human ortholog. The second group of mutants consisted of fish that had been injected with a protein trap transposon system that truncates the expected protein products as well as sorting and imaging through fluorescent expression patterns *in vivo*. Included here, we have curated small group of mutants where the GBT integrated into a mitochondrial gene of interest.

In recent years zebrafish have been used to understand the pathophysiology of human mitochondrial disorders such as cardiovascular, multisystemic, neurological, and erythropoiesis, to name a few. Studies for loss of function have been carried out to model left ventricular non-compaction, a genetically heterogeneous cardiomyopathy caused by mutations in nicotinamide nucleotide transhydrogenase (*NNT*) gene [98,99]. Transiently suppressing the levels of *nnt* gene did mirror the symptoms, such as early ventricular malformation and contractility defects, observed in human patients [99]. Transient suppression of *ndufb11* protein in zebrafish results in cardiac anomalies, confirming the role of *NDUFB11* in histiocytoid cardiomyopathy pathogenesis [41].

Parkinson disease (PD) is another example where *parkin* and *pink* knockdown larvae display mitochondrial fragmentation accompanied by accumulation of oxidative species, mimicking the phenotype observed in PD patients. Loss of dendritic arborization and peripheral neurons were also observed in the knockdown animals [72,100–103]. The zebrafish model of Dravet syndrome shows altered metabolism associated with the pathophysiology [104]. Downregulation of the sodium channel, voltage-gated, type I like, alpha b (*scnl1ab*) in zebrafish leads to mitochondrial structural abnormalities accompanied by accumulation of oxidative species [104–106].

Sideroblastic anemia, caused by mutations in mitoferrin (*mfrn1*) gene, has been successfully modelled in zebrafish [107]. The elucidation of its role in zebrafish as iron importer does shed some light on iron related defects in progression of anemia, accompanied by mitochondrial dysfunction. Zebrafish erythrocytes are nucleated and possess organelles such as mitochondria [108] in their cytoplasm, making them an ideal model organism to study ribosomopathies and hematological malignancies [109–114]. Mitochondria in erythrocytes may offer valuable insight into novel biological pathways resident in mitochondria.

Transient infantile liver failure and mitochondrial deafness, caused by mutations in tRNA 5-methylaminomethyl-2-thiouridylate methyltransferase (*TRMU*) gene [115] have been phenocopied in zebrafish [116]. Zebrafish *trmu* mutants exhibit abnormal mitochondrial morphology, deficient oxidative phosphorylation, defects in hair cells and decreases in the steady state levels of mt-tRNAs, resembling the observations made in patients suffering from this syndrome [116].

Taking cues from these studies, many of the genes selected here were prioritized on the basis of the mutations and clinical phenotype reported in patients. Detailed phenotypic analyses will be described in subsequent studies. The zebrafish mutants encode for genes that have been involved in myriad of clinical phenotypes such as encephalopathy, nephrotic syndrome, intellectual disability, psychomotor developmental delay, respiratory chain deficiency, lactic acidosis, iron overload, microcephaly, hypertrophic cardiomyopathy, etc. The superficial investigations of the tissue-specific phenotypes associated with, for example, housekeeping nuclear encoded proteins, have to be refined with organelle level investigations. The repository of mutants encoding for proteins involved in respiratory chain biogenesis and assembly offers the potential to understand the moonlighting role of these proteins. Mutations in these proteins are associated with organ-specific phenotypes such as neurological, paraganglioma, cardiovascular etc. [41,117]. Extensive energy demand by these tissues is one possible explanation for the progression of these manifestations; however, the role of these proteins in organ-specific cellular niches or homeostasis is also a possibility. These models can help to decipher the role of these proteins in the mitochondrial interactome, when studied *in vivo*. This underpins the utility of this MMC collection in deciphering the role of mitochondrial proteins in tissue-specific biological pathways.

Zebrafish are also amenable to small molecule mediated pharmacological modelling of mitochondrial phenotypes. To test hypoxia as a potential protective therapy in mitochondrial disorders, Von Hippel-Lindau (*vhl*) null mutant zebrafish, when treated with antimycin mediated mitochondrial insult, exhibited improved survival. In addition, FG-4592 was found to improve survival in response to respiratory chain inhibition, possibly due to increase in hypoxia response [118]. A series of ETC complex specific pharmacological inhibitors such as rotenone (complex I), azide (complex IV), oligomycin (complex V) and chloramphenicol (mitochondrial protein translation) have been used to model respiratory chain dysfunction in zebrafish [78,119,120]. Zebrafish larvae display a series of phenotypes, such as developmental arrest, when treated with rotenone. Treatment with azide induced decreased heart rate, loss of motor function, inability to respond to tactical stimulation, neurological damage and mortality. Organ specific manifestations such as neurological and behavioral dysfunction have been reported in 6-7 days post fertilization (dpf) zebrafish larvae upon administration of a titred drug concentration [120]. These observations tilt the scales in favor of zebrafish for modeling mitochondrial clinical manifestations. Gaining insights from these studies, our compendium of mutants provides an excellent resource to test a large number of biological and chemical modulators as potential therapies for mitochondrial disorders for which treatment remains elusive.

Zebrafish offer many unique advantages for *in vivo* imaging experiments, as compared to their mammalian counterparts. A classical study corroborating this advantage is *in vivo* imaging of mitochondrial transport in a single axon, as demonstrated by Yang and his colleagues [121]. Seok and his colleagues generated a transgenic zebrafish line, expressing green fluorescent protein fused to mitochondrial localization sequence from cytochrome c oxidase [71]. Mitochondrial function is often measured by various parameters such as estimation of membrane potential, mitochondrial superoxide species and energy production. Superoxide activity and membrane potential in zebrafish has been measured by employing the use of cell permeable chemical probes such as MitoSox [122] and Dihydrorhodamine 123 (DH123) [73], respectively. Zebrafish transgenic lines expressing genetically encoded calcium and oxidation indicators, have also served as an excellent model to measure calcium homeostasis and oxidative status *in vivo* in models of mechanosensory hair cell damage and death [123]. Constructs such as Mitotimer allow the measurement of mitochondrial turnover, transport, and changes in the redox history of mitochondria during organogenesis events in zebrafish embryos [124]. This transgenic model enables a sweeping picture of the mitochondrial network, helping in studying cellular processes such as mitophagy and apoptosis.

Zebrafish have also provided novel mechanistic insights to interrogate the mitochondrial dynamics, to be called as “*in-vivo* life cycle of mitochondria” in healthy and diseased conditions [124]. One such example is that of Mitofish [74], that recapitulates mitochondrial network biogenesis, unraveling the role of the organelle in cell-type specific niches of different organ systems. Mitofish is a transgenic zebrafish line that fluorescently labels the mitochondria in the neurons, enabling non-invasive *in vivo* observation. Advancement in adaptive optics and lattice light sheet microscopy have empowered researchers to look at vibrant and colorful images of organelles in zebrafish. These technologies have been applied in investigating the organellar dynamics in brain during early development and in the eye of an adult zebrafish [125]. These MMMC mutants provide an excellent platform to study mitochondrial remodeling and homeostasis in the progression of various clinical disorders by mapping the mitochondrial wiring of the cell.

These zebrafish mutants have been cryopreserved for the purpose of sharing the lines as a resource with the field for expanding the study of mitochondrial genetics and biology. In addition, our template for the creation of mitochondrial mutants in zebrafish should enable the creation of higher animal models of specific variants of interest. For the purpose of accelerating this research, all lines in this project have had their information deposited with ZIRC, and all of the lines have been made available via zfishbook (www.zfishbook.com) [83].

The goal of this study is to disseminate the use of zebrafish as a *noblesse oblige* in the field of mitochondrial biology and medicine, paving way for the development of novel insight for diagnostic and therapeutic strategies. Ultimately, using this collection of mutants we hope to unravel a small part of the mystery that shrouds one of the most crucial organelles in the cell. We hope that this MMMC mutant collection and the primer we provide for adding to it, will help to usher in a new mitochondrial research using the zebrafish.

## Acknowledgements

The authors thank Dr. Eric Schon and Dr. Vamsi Mootha for helping select the gene set targeted in this collection, and the following who contributed to the generation of these new zebrafish lines including Alexandra Cook, Roberto Lopez Cervera, and the Mayo Clinic Zebrafish Facility staff. Gabriel Martinez Galvez helped with paper illustration design. This work was sponsored by the Mayo Foundation, a gift by the Marriott Foundation and by NIH grants DK84567, GM63904 and HG006431 to SCE.

## Author Contributions

The paper was written by AS, MW, ZWJ and SCE. Experiments were executed by JMC, MW, HA, NI, MDU, JDB, RMH, WL, YD and TLP with experimental guidance from XX, KJC and SCE. Data analysis was completed by JMC, NI, JDB, MDU, RMH, MW, WL, YD and AS.

