## Supplementary table 1 for "A primer genetic toolkit for exploring mitochondrial biology and disease using zebrafish"

### Supplementary File

**Supplementary Table 1: List of mitochondrial respiratory chain complex genes:** The current resource list summarizes information on human gene, gene family description, zebrafish ortholog, mouse ortholog, relevant clinical phenotypes and diseases, and OMIM ID. Blue = Nuclear genome encoded mitochondrial proteins, Mustard = Mitochondrial genome encoded proteins (Keywords: Chr- Chromosome; OMIM ID: Online Mendelian Inheritance in Man ID; NA-not available).

| **Complex 1- NADH ubiquinone oxidoreductase subunits** | | | | | | | | | | |
|---|---|---|---|---|---|---|---|---|---|---|
| **Approved symbol** | **Approved name** | **Chr** | **Gene family description** | **Zebrafish orthologs** | **Zebrafish chr** | **Mouse orthologs** | **Mouse chr** | **Clinical manifestations observed in humans** | **Disease** | **OMIM ID** |
| *NDUFS1* | NADH:ubiquinone oxidoreductase core subunit S1 | 2 | NADH:ubiquinone oxidoreductase core subunits | *ndufs1* | 6 | *Ndufs1* | 1 | Growth retardation, Psychomotor retardation with hypotonia, microcephaly, nystagmus, bilateral optic atrophy, leukodystrophy, leukoencephalopathy hyperlactatemia and hyperlactatorachia | Mitochondrial complex-I deficiency | 157655 |
| *NDUFV1* | NADH:ubiquinone oxidoreductase core subunit V1 | 11 | NADH:ubiquinone oxidoreductase core subunits | *ndufv1* | 19 | *Ndufv1* | 19 | Encephalopathy, ataxia, bilateral ptosis, and ophthalmoplegia, lactic acidosis, leukodystrophy | Mitochondrial complex-I deficiency | 161015 |
| *NDUFS2* | NADH:ubiquinone oxidoreductase core subunit S2 | 1 | NADH:ubiquinone oxidoreductase core subunits | *ndufs2* | 7 | *Ndufs2* | 1 | Nystagmus, bilateral optic atrophy, hypotonia, axial hyperlactemia, neurological regression, optic atrophy, epilepsy, psychomotor delay, dystonia and leukoencephalopathy | Mitochondrial complex-I deficiency | 602985 |
| *NDUFS3* | NADH:ubiquinone oxidoreductase core subunit S3 | 11 | NADH:ubiquinone oxidoreductase core subunits | *ndufs3* | 7 | *Ndufs3* | 2 | Mitochondrial encephalopathy, mitochondrial myopathy, developmental delay, muscular hypotonia and lactic acidosis | Leigh syndrome; Mitochondrial complex-I deficiency | 603846 |

| Approved symbol | Approved name | Chr | Gene family description | Zebrafish orthologs | Zebrafish chr | Mouse orthologs | Mouse chr | Clinical manifestations observed in humans | Disease | OMIM ID |
|---|---|---|---|---|---|---|---|---|---|---|
| *NDUFV2* | NADH:ubiquinone oxidoreductase core subunit V2 | 18 | NADH:ubiquinone oxidoreductase core subunits | *ndufv2* | 2 | *Ndufv2* | 17 | Hypertrophic cardiomyopathy, hypotonia, growth retardation, hyperlactatemia, encephalomyopathy, epilepsy, progressive spasticity and nystagmus | Mitochondrial complex-I deficiency | 600532 |
| *NDUFS7* | NADH:ubiquinone oxidoreductase core subunit S7 | 19 | NADH:ubiquinone oxidoreductase core subunits | *ndufs7* | 11 | *Ndufs7* | 10 | Hemorrhagic diathesis, ataxia, muscular hypotonia, nystagmus, hepatic and renal failure, multifocal hyperechogenicity, hypotonia neurological regression and muscular weakness | Leigh syndrome | 601825 |
| *NDUFS8* | NADH:ubiquinone oxidoreductase core subunit S8 | 11 | NADH:ubiquinone oxidoreductase core subunits | *ndufs8a*<br>*ndufs8b* | 1<br>14 | *Ndufs8* | 19 | Muscular hypotonia, delayed motor milestones, mild ptosis, muscle weakness, dysarthia, ataxic gait, nystagmus and balance impairment | Leigh syndrome; Mitochondrial complex-I deficiency | 602141 |
| *MT-ND1* | Mitochondrially encoded NADH:ubiquinone oxidoreductase core subunit 1 | MT | NADH:ubiquinone oxidoreductase core subunits | *mt-nd1* | MT | *mt-Nd1* | MT | Optic atrophy, Alzheimer, sudden infant death, exercise intolerance, dystonia, lactic acidosis and sensorineural deafness | Mitochondrial complex-1 deficiency; Leber optic atrophy; MELAS; Sudden death infant syndrome; Dystonia | 516000 |
| *MT-ND2* | Mitochondrially encoded NADH:ubiquinone oxidoreductase core subunit 2 | MT | NADH:ubiquinone oxidoreductase core subunits | *mt-nd2* | MT | *mt-Nd2* | MT | Optic atrophy, exercise intolerance, encephalomyopathy and lactic acidosis | Mitochondrial complex-1 deficiency; Leber optic atrophy; Leigh syndrome | 516001 |
| *MT-ND3* | Mitochondrially encoded NADH:ubiquinone oxidoreductase core subunit 3 | MT | NADH:ubiquinone oxidoreductase core subunits | *mt-nd3* | MT | *mt-Nd3* | MT | Myoclonus, seizures, cognitive decline, ataxia, peripheral neuropathy, eye movement abnormalities, encephalopathy and optic atrophy | Mitochondrial complex-1 deficiency; Leigh syndrome | 516002 |
| *MT-ND4* | Mitochondrially encoded NADH:ubiquinone oxidoreductase core subunit 4 | MT | NADH:ubiquinone oxidoreductase core subunits | *mt-nd4* | MT | *mt-Nd4* | MT | Optic atrophy, cardiomyopathy, dystonia, intermittent migraines, sensorineural hearing loss, bilateral cataracts, grand mal seizures, stroke-like episodes, lactic acidosis, and ragged-red muscle fibers | Leber optic atrophy; MELAS; Dystonia | 516003 |
| *MT-ND4L* | Mitochondrially encoded NADH:ubiquinone oxidoreductase core subunit 4L | MT | NADH:ubiquinone oxidoreductase core subunits | *mt-nd4l* | MT | *mt-Nd4l* | MT | Optic atrophy and colorectal cancer | Leber optic atrophy; Colorectal cancer | 516004 |

| Approved symbol | Approved name | Chr | Gene family description | Zebrafish orthologs | Zebrafish chr | Mouse orthologs | Mouse chr | Clinical manifestations observed in humans | Disease | OMIM ID |
|---|---|---|---|---|---|---|---|---|---|---|
| *MT-ND5* | Mitochondrially encoded NADH:ubiquinone oxidoreductase core subunit 5 | MT | NADH:ubiquinone oxidoreductase core subunits | *mt-nd5* | MT | *mt-Nd5* | MT | Ataxia, facial weakness, impaired hearing, ophthalmoplegia, exercise intolerance, developmental delay, myoclonus and neurological dysfunction | Mitochondrial complex-1 deficiency; Leber optic atrophy; Leigh syndrome; MELAS | 516005 |
| *MT-ND6* | Mitochondrially encoded NADH:ubiquinone oxidoreductase core subunit 6 | MT | NADH:ubiquinone oxidoreductase core subunits | *mt-nd6* | MT | *mt-Nd6* | MT | Optic atrophy, dystonia, anarthria, spasticity, motor retardation, motor retardation, mild encephalopathy and oncocytoma | Mitochondrial complex-1 deficiency; Leber optic atrophy; Leigh syndrome; MELAS; Oncocytoma | 516006 |

**Complex 1- NADH ubiquinone oxidoreductase assembly**

| Approved symbol | Approved name | Chr | Gene family description | Zebrafish orthologs | Zebrafish chr | Mouse orthologs | Mouse chr | Clinical manifestations observed in humans | Disease | OMIM ID |
|---|---|---|---|---|---|---|---|---|---|---|
| *NDUFA1* | NADH:ubiquinone oxidoreductase subunit A1 | X | NADH:ubiquinone oxidoreductase supernumerary subunits | *ndufa1* | 14 | *Ndufa1* | X | Hypotonia, nystagmus, epilepsy, lactic acidosis, neurodegeneration, developmental delay and ataxia, hypotonia, nystagmus, epilepsy, lactic acidosis, neurodegeneration, developmental delay and ataxia | Mitochondrial complex-1 deficiency | 300078 |
| *NDUFA10* | NADH:ubiquinone oxidoreductase subunit A10 | 2 | NADH:ubiquinone oxidoreductase supernumerary subunits | *ndufa10* | 9 | *Ndufa10* | 1 | Hypotonia, muscular hypotrophy, gait ataxia, growth retardation and lactic acidosis | Leigh syndrome | 603835 |
| *NDUFA11* | NADH:ubiquinone oxidoreductase subunit A11 | 19 | NADH:ubiquinone oxidoreductase supernumerary subunits | *ndufa11* | 11 | *Ndufa11* | 17 | Hyperlactatemia, impaired psychomotor development, microcephaly, hypotonia, muscle weakness and nystagmus | Mitochondrial complex-I deficiency | 612638 |
| *NDUFA12* | NADH:ubiquinone oxidoreductase subunit A12 | 12 | NADH:ubiquinone oxidoreductase supernumerary subunits | *ndufa12* | 4 | *Ndufa12* | 10 | Delayed early motor development, balance impairment, scoliosis, dystonia, growth retardation, muscular atrophy and hypotonia | Mitochondrial complex-I deficiency | 614530 |
| *NDUFA13* | NADH:ubiquinone oxidoreductase subunit A13 | 19 | NADH:ubiquinone oxidoreductase supernumerary subunits | *ndufa13* | 1 | *Ndufa13* | 8 | Papillary thyroid carcinoma | Thyroid carcinoma | 609435 |
| *NDUFA2* | NADH:ubiquinone oxidoreductase subunit A2 | 5 | NADH:ubiquinone oxidoreductase supernumerary subunits | *ndufa2* | 21 | *Ndufa2* | 18 | Developmental delay, cardiomyopathy, cerebral atrophy and encephalopathy | Leigh syndrome | 602137 |
| *NDUFA3* | NADH:ubiquinone oxidoreductase subunit A3 | 19 | NADH:ubiquinone oxidoreductase supernumerary subunits | *ndufa3* | 16 | *Ndufa3* | 7 | -na- | -na- | 603832 |
| *NDUFA5* | NADH:ubiquinone oxidoreductase subunit A5 | 7 | NADH:ubiquinone oxidoreductase supernumerary subunits | *ndufa5* | 4 | *Ndufa5* | 6 | -na- | -na- | 601677 |
| *NDUFA6* | NADH:ubiquinone oxidoreductase subunit A6 | 22 | NADH:ubiquinone oxidoreductase supernumerary subunits | *ndufa6* | 3 | *Ndufa6* | 15 | -na- | -na- | 602138 |

| Approved symbol | Approved name | Chr | Gene family description | Zebrafish orthologs | Zebrafish chr | Mouse orthologs | Mouse chr | Clinical manifestations observed in humans | Disease | OMIM ID |
|---|---|---|---|---|---|---|---|---|---|---|
| *NDUFA7* | NADH:ubiquinone oxidoreductase subunit A7 | 19 | NADH:ubiquinone oxidoreductase supernumerary subunits | *ndufa7* | 8 | *Ndufa7* | 17 | -na- | -na- | 602139 |
| *NDUFA8* | NADH:ubiquinone oxidoreductase subunit A8 | 9 | NADH:ubiquinone oxidoreductase supernumerary subunits | *ndufa8* | 10 | *Ndufa8* | 2 | Neonatal hypotonia, dysmorphic features, epilepsy and lactic acidosis in plasma | Mitochondrial complex-1 deficiency | 603359 |
| *NDUFA9* | NADH:ubiquinone oxidoreductase subunit A9 | 12 | NADH:ubiquinone oxidoreductase supernumerary subunits | *ndufa9a*<br>*ndufa9b* | 25<br>10 | *Ndufa9* | 6 | Metabolic acidosis, dystonia, dysphagia, speech disturbance and dysarthria | Leigh syndrome | 603834 |
| *NDUFAB1* | NADH:ubiquinone oxidoreductase subunit AB1 | 16 | NADH:ubiquinone oxidoreductase supernumerary subunits | *ndufab1a*<br>*ndufab1b* | 1<br>3 | *Ndufab1* | 7 | -na- | -na- | 603836 |
| *NDUFB1* | NADH:ubiquinone oxidoreductase subunit B1 | 14 | NADH:ubiquinone oxidoreductase supernumerary subunits | *ndufb1* | 20 | *Ndufb1* | 12 | -na- | -na- | 603837 |
| *NDUFB10* | NADH:ubiquinone oxidoreductase subunit B10 | 16 | NADH:ubiquinone oxidoreductase supernumerary subunits | *ndufb10* | 3 | *Ndufb10* | 17 | -na- | -na- | 603843 |
| *NDUFB11* | NADH:ubiquinone oxidoreductase subunit B11 | X | NADH:ubiquinone oxidoreductase supernumerary subunits | *ndufb11* | 8 | *Ndufb11* | X | Histiocytoid cardiomyopathy and microphthalmia with linear skin defects syndrome and axial hypotonia | Linear skin defects with multiple congenital anomalies; Mitochondrial complex-I deficiency | 300403 |
| *NDUFB2* | NADH:ubiquinone oxidoreductase subunit B2 | 7 | NADH:ubiquinone oxidoreductase supernumerary subunits | *ndufb2* | 4 | *Ndufb2* | 6 | -na- | -na- | 603838 |
| *NDUFB3* | NADH:ubiquinone oxidoreductase subunit B3 | 2 | NADH:ubiquinone oxidoreductase supernumerary subunits | *ndufb3* | 9 | *Ndufb3* | 1 | Hypoglycemia, intrauterine growth restriction, hypertrophic cardiomyopathy, failure to thrive, muscular hypotonia and lactic acidosis | Mitochondrial complex-I deficiency | 603839 |
| *NDUFB4* | NADH:ubiquinone oxidoreductase subunit B4 | 3 | NADH:ubiquinone oxidoreductase supernumerary subunits | *ndufb4* | 24 | *Ndufb4* | 16 | -na- | -na- | 603840 |
| *NDUFB5* | NADH:ubiquinone oxidoreductase subunit B5 | 3 | NADH:ubiquinone oxidoreductase supernumerary subunits | *ndufb5* | 6 | *Ndufb5* | 3 | -na- | -na- | 603841 |
| *NDUFB6* | NADH:ubiquinone oxidoreductase subunit B6 | 9 | NADH:ubiquinone oxidoreductase supernumerary subunits | *ndufb6* | 1 | *Ndufb6* | 4 | -na- | -na- | 603322 |
| *NDUFB7* | NADH:ubiquinone oxidoreductase subunit B7 | 19 | NADH:ubiquinone oxidoreductase supernumerary subunits | *ndufb7* | 1 | *Ndufb7* | 8 | -na- | -na- | 603842 |
| *NDUFB8* | NADH:ubiquinone oxidoreductase subunit B8 | 10 | NADH:ubiquinone oxidoreductase supernumerary subunits | *ndufb8* | 12 | *Ndufb8* | 19 | Hypotonia, failure to thrive, delayed psychomotor development, seizures and lactic acidosis | Mitochondrial complex-I deficiency | 602140 |
| *NDUFB9* | NADH:ubiquinone oxidoreductase subunit B9 | 8 | NADH:ubiquinone oxidoreductase supernumerary subunits | *ndufb9* | 16 | *Ndufb9* | 15 | Progressive hypotonia | Mitochondrial complex-I deficiency | 601445 |

| Approved symbol | Approved name | Chr | Gene family description | Zebrafish orthologs | Zebrafish chr | Mouse orthologs | Mouse chr | Clinical manifestations observed in humans | Disease | OMIM ID |
|---|---|---|---|---|---|---|---|---|---|---|
| *NDUFC1* | NADH:ubiquinone oxidoreductase subunit C1 | 4 | NADH:ubiquinone oxidoreductase supernumerary subunits | *si:ch211-235e9.6* | 14 | *Ndufc1* | 3 | -na- | -na- | 603844 |
| *NDUFC2* | NADH:ubiquinone oxidoreductase subunit C2 | 11 | NADH:ubiquinone oxidoreductase supernumerary subunits | *ndufc2* | 15 | *Ndufc2* | 7 | -na- | -na- | 603845 |
| *NDUFV3* | NADH:ubiquinone oxidoreductase subunit V3 | 21 | NADH:ubiquinone oxidoreductase supernumerary subunits | *ndufv3* | 9 | *Ndufv3* | 17 | -na- | -na- | 602184 |
| *NDUFS4* | NADH:ubiquinone oxidoreductase subunit S4 | 5 | NADH:ubiquinone oxidoreductase supernumerary subunits | *ndufs4* | 5 | *Ndufs4* | 13 | Mitochondrial complex I deficiency | Mitochondrial complex-I deficiency | 602694 |
| *NDUFS5* | NADH:ubiquinone oxidoreductase subunit S5 | 1 | NADH:ubiquinone oxidoreductase supernumerary subunits | *ndufs5* | 19 | *Ndufs5* | 4 | -na- | -na- | 603847 |
| *NDUFS6* | NADH:ubiquinone oxidoreductase subunit S6 | 5 | NADH:ubiquinone oxidoreductase supernumerary subunits | *ndufs6* | 19 | *Ndufs6* | 13 | Encephalopathy, lactic acidosis, metabolic acidosis, hypotonia and infant death | Mitochondrial complex-I deficiency | 603848 |
| **Complex 2- Succinate dehydrogenase subunits** | | | | | | | | | | |
| **Approved symbol** | **Approved name** | **Chr** | **Gene family description** | **Zebrafish orthologs** | **Zebrafish chr** | **Mouse orthologs** | **Mouse chr** | **Clinical manifestations observed in humans** | **Disease** | **OMIM ID** |
| *SDHA* | Succinate dehydrogenase complex flavoprotein subunit A | 5 | Mitochondrial complex II: succinate dehydrogenase subunits | *sdha* | 19 | *Sdha* | 13 | Dyspnea, cardiomegaly, cardiomyopathy, nystagmus, hypotonia, gastrointestinal stromal tumors, paragangliomas, pheochromocytoma, psychomotor regression, severe hyperandrogenism | Mitochondrial complex-II deficiency; Cardiomyopathy; Leigh syndrome; Paraganglioma | 600857 |
| *SDHB* | Succinate dehydrogenase complex iron sulfur subunit B | 1 | Mitochondrial complex II: succinate dehydrogenase subunits | *sdhb* | 22 | *Sdhb* | 4 | Pheochromocytoma, paraganglioma, dysphonia, dysphagia and cardiomyopathy | Cowden syndrome; Gastrointestinal stromal tumor; Paraganglioma; Gastric stromal sarcoma; Pheochromocyto--ma | 185470 |
| *SDHC* | Succinate dehydrogenase complex subunit C | 1 | Mitochondrial complex II: succinate dehydrogenase subunits | *sdhc* | 2 | *Sdhc* | 1 | Paraganglioma, prolactinoma and pituitary gangliocytoma | Gastrointestinal stromal tumor; Paraganglioma and gastric stromal sarcoma | 602413 |
| *SDHD* | succinate dehydrogenase complex subunit D | 11 | Mitochondrial complex II: succinate dehydrogenase subunits | *sdhda*<br>*sdhdb* | 5<br>15 | *Sdhd* | 9 | Primary renal paragangliomas, renal neoplasia, pheochromocytomas, head-neck paragangliomas and sensorineural hearing loss | Mitochondrial complex-II deficiency; Cowden syndrome 3; Paraganglioma 1 with or without deafness; Gastric stromal sarcoma; Pheochromocytoma; Merkel cell carcinoma somatic; Carcinoid intestinal tumors | 602690 |

| Complex 3- Ubiquinol cytochrome c reductase complex subunits | | | | | | | | | | |
|---|---|---|---|---|---|---|---|---|---|---|
| **Approved symbol** | **Approved name** | **Chr** | **Gene family description** | **Zebrafish orthologs** | **Zebrafish chr** | **Mouse orthologs** | **Mouse chr** | **Clinical manifestations observed in humans** | **Disease** | **OMIM ID** |
| *CYC1* | cytochrome c1 | 8 | Mitochondrial complex III: ubiquinol-cytochrome c reductase complex subunits | *cyc1* | 6 | *Cyc1* | 15 | Ketoacidosis, hyperlactatemia, hyperammonemia, mild growth retardation and congenital left ptosis, hyperlactatemia and insulin-responsive hyperglycemia | Mitochondrial complex III deficiency | 123980 |
| *UQCR10* | ubiquinol-cytochrome c reductase, complex III subunit X | 22 | Mitochondrial complex III: ubiquinol-cytochrome c reductase complex subunits | *uqcr10* | 21 | *Uqcr10* | 11 | -na- | -na- | 610843 |
| *UQCR11* | ubiquinol-cytochrome c reductase, complex III subunit XI | 19 | Mitochondrial complex III: ubiquinol-cytochrome c reductase complex subunits | *uqcr11* | 2 | *Uqcr11* | 10 | -na- | -na- | 609711 |
| *UQCRB* | ubiquinol-cytochrome c reductase binding protein | 8 | Mitochondrial complex III: ubiquinol-cytochrome c reductase complex subunits | *uqcrb* | 19 | *Uqcrb* | 13 | Hepatomegaly, hypoglycemia, metabolic acidosis, hyperlactatemia and hyperlactatemia | Mitochondrial complex III deficiency | 191330 |
| *UQCRC1* | ubiquinol-cytochrome c reductase core protein 1 | 3 | Mitochondrial complex III: ubiquinol-cytochrome c reductase complex subunits | *uqcrc1* | 6 | *Uqcrc1* | 9 | -na- | -na- | 191328 |
| *UQCRC2* | ubiquinol-cytochrome c reductase core protein 2 | 16 | Mitochondrial complex III: ubiquinol-cytochrome c reductase complex subunits | *uqcrc2a*<br>*uqcrc2b* | 3<br>12 | *Uqcrc2* | 7 | Lactic acidosis, hypoglycemia, ketosis and hyperammonemia | Mitochondrial complex III deficiency | 191329 |
| *UQCRFS1* | ubiquinol-cytochrome c reductase, Rieske iron-sulfur polypeptide 1 | 19 | Mitochondrial complex III: ubiquinol-cytochrome c reductase complex subunits | *uqcrfs1* | 7 | *Uqcrfs1* | 13 | -na- | -na- | 191327 |
| *UQCRH* | ubiquinol-cytochrome c reductase hinge protein | 1 | Mitochondrial complex III: ubiquinol-cytochrome c reductase complex subunits | *uqcrh* | 6 | *Uqcrh* | 4 | -na- | -na- | 613844 |
| *UQCRQ* | ubiquinol-cytochrome c reductase complex III subunit VII | 5 | Mitochondrial complex III: ubiquinol-cytochrome c reductase complex subunits | *uqcrq* | 14 | *Uqcrq* | 11 | Severe psychomotor retardation and extrapyramidal signs, dystonia, athetosis and ataxia, mild axial hypotonia and marked global dementia | Mitochondrial complex III deficiency | 612080 |

| Approved symbol | Approved name | Chr | Gene family description | Zebrafish orthologs | Zebrafish chr | Mouse orthologs | Mouse chr | Clinical manifestations observed in humans | Disease | OMIM ID |
|---|---|---|---|---|---|---|---|---|---|---|
| *MT-CYB* | mitochondrially encoded cytochrome b | MT | Mitochondrial complex III: ubiquinol-cytochrome c reductase complex subunits | *mt-cyb* | MT | *mt-Cyb* | MT | Optic atrophy, colorectal cancer, exercise intolerance, encephalomyopathy, lactic acidosis, cardiomyopathy and septo-optic dysplasia | Leber optic atrophy; Colorectal cancer; Exercise intolerance; Encephalomyop-ahty; Infantile histiocytoid cardiomyopathy; Septo-optic dysplasia; MELAS | 516020 |
| **Complex 4- Cytochrome c oxidase subunits** | | | | | | | | | | |
| **Approved symbol** | **Approved name** | **Chr** | **Gene family description** | **Zebrafish orthologs** | **Zebrafish chr** | **Mouse orthologs** | **Mouse chr** | **Clinical manifestations observed in humans** | **Disease** | **OMIM ID** |
| *COX4I1* | cytochrome c oxidase subunit 4I1 | 16 | Mitochondrial complex IV: cytochrome c oxidase subunits | *cox4i1* | 18 | *Cox4i1* | 8 | -na- | -na- | 123864 |
| *COX4I2* | cytochrome c oxidase subunit 4I2 | 20 | Mitochondrial complex IV: cytochrome c oxidase subunits | *cox4i2* | 23 | *Cox4i2* | 2 | Congenital exocrine pancreatic insufficiency, dyserythropoeitic anemia, calvarial hyperostosis and intellectual disability | Exocrine pancreatic insufficiency, dyserythropoietic anemia, and calvarial hyperostosis | 607976 |
| *COX5A* | cytochrome c oxidase subunit 5A | 15 | Mitochondrial complex IV: cytochrome c oxidase subunits | *cox5aa*<br>*cox5ab* | 25<br>18 | *Cox5a* | 9 | -na- | -na- | 603773 |
| *COX5B* | cytochrome c oxidase subunit 5B | 2 | Mitochondrial complex IV: cytochrome c oxidase subunits | *cox5b* | 10 | *Cox5b* | 1 | -na- | -na- | 123866 |
| *COX6A1* | cytochrome c oxidase subunit 6A1 | 12 | Mitochondrial complex IV: cytochrome c oxidase subunits | *cox6a1* | 8 | *Cox6a1* | 5 | Axonal hereditary motor and sensory neuropathy | Charcot-Marie-Tooth disease, recessive intermediate | 602072 |
| *COX6A2* | cytochrome c oxidase subunit 6A2 | 16 | Mitochondrial complex IV: cytochrome c oxidase subunits | *cox6a2* | 3 | *Cox6a2* | 7 | -na- | -na- | 6002009 |
| *COX6B1* | cytochrome c oxidase subunit 6B1 | 19 | Mitochondrial complex IV: cytochrome c oxidase subunits | *cox6b1* | 24 | *Cox6b1* | 7 | Leukodystrophy, muscle weakness, cognitive deterioration, visual loss, metabolic acidosis and lactic acidosis | Mitochondrial complex IV deficiency | 124089 |
| *COX6B2* | cytochrome c oxidase subunit 6B2 | 19 | Mitochondrial complex IV: cytochrome c oxidase subunits | *cox6b2* | 16 | *Cox6b2* | 7 | -na- | -na- | 618127 |
| *COX6C* | cytochrome c oxidase subunit 6C | 8 | Mitochondrial complex IV: cytochrome c oxidase subunits | *cox6c* | 6 | *Cox6c* | 15 | -na- | -na- | 124090 |
| *COX7A1* | cytochrome c oxidase subunit 7A1 | 19 | Mitochondrial complex IV: cytochrome c oxidase subunits | *cox7a1* | 15 | *Cox7a1* | 7 | -na- | -na- | 123995 |

| Approved symbol | Approved name | Chr | Gene family description | Zebrafish orthologs | Zebrafish chr | Mouse orthologs | Mouse chr | Clinical manifestations observed in humans | Disease | OMIM ID |
|---|---|---|---|---|---|---|---|---|---|---|
| *COX7A2* | cytochrome c oxidase subunit 7A2 | 6 | Mitochondrial complex IV: cytochrome c oxidase subunits | *cox7a2a* *cox7a2b* | 17 20 | *Cox7a2* | 9 | -na- | -na- | 605771 |
| *COX7B* | cytochrome c oxidase subunit 7B | X | Mitochondrial complex IV: cytochrome c oxidase subunits | *cox7b* | 21 | *Cox7b* | X | Linear skin lesions, facial dysmorphisms, short stature, microcephaly, poor vision and intellectual disabilities | Linear skin defects with multiple congenital anomalies | 300885 |
| *COX7B2* | cytochrome c oxidase subunit 7B2 | 4 | Mitochondrial complex IV: cytochrome c oxidase subunits | -na- | -na- | *Cox7b2* | 5 | -na- | -na- | 609811 |
| *COX7C* | cytochrome c oxidase subunit 7C | 5 | Mitochondrial complex IV: cytochrome c oxidase subunits | *cox7c* | 5 | *Cox7c* | 13 | -na- | -na- | 603774 |
| *COX8A* | cytochrome c oxidase subunit 8A | 11 | Mitochondrial complex IV: cytochrome c oxidase subunits | *cox8a* | 21 | *Cox8a* | 19 | Developmental delay, seizures, lactic acidosis, psychomotor psychomotor retardation, a short stature, microcephalus, exophthalmos, proximal muscular hypotonia, distal spasticity, pigmentary retinopathy and metabolic encephalopathy | Mitochondrial complex IV deficiency | 123870 |
| *COX8C* | cytochrome c oxidase subunit 8C | 14 | Mitochondrial complex IV: cytochrome c oxidase subunits | -na- | -na- | *Cox8c* | 12 | -na- | -na- | 616855 |
| *MT-CO1* | mitochondrially encoded cytochrome c oxidase I | MT | Mitochondrial complex IV: cytochrome c oxidase subunits | *mt-co1* | MT | *mt-Co1* | MT | Optic atrophy, sensorineural deafness, sideroblastic anemia, ataxia, myoclonic epilepsy, mental retardation, colorectal cancer and myoglobinuria | Leber optic atrophy; Acquired idiopathic sideroblastic anemia; Cytochrome c oxidase deficiency; Colorectal cancer; Myoglobinuria | 516030 |
| *MT-CO2* | mitochondrially encoded cytochrome c oxidase II | MT | Mitochondrial complex IV: cytochrome c oxidase subunits | *mt-co2* | MT | *mt-Co2* | MT | Optic atrophy, ataxia, lactic acidosis and colorectal cancer | Cytochrome c oxidase deficiency | 516040 |
| *MT-CO3* | mitochondrially encoded cytochrome c oxidase III | MT | Mitochondrial complex IV: cytochrome c oxidase subunits | *mt-co3* | MT | *mt-Co3* | MT | Optic atrophy, seizures, lactic acidosis, myopathy, exercise intolerance, delayed growth and myoglobinuria | Leber optic atrophy; Mitochondrial complex-IV deficiency | 516050 |

**Complex 5- ATP synthase subunits**

| Approved symbol | Approved name | Chr | Gene family description | Zebrafish orthologs | Zebrafish chr | Mouse orthologs | Mouse chr | Clinical manifestations observed in humans | Disease | OMIM ID |
|---|---|---|---|---|---|---|---|---|---|---|
| *ATP5F1A* | ATP synthase F1 subunit alpha | 18 | Mitochondrial complex V: ATP synthase subunits | *atp5f1a* | 21 | *Atp5f1a* | 18 | Encephalopathy | Mitochondrial complex (ATP synthase) deficiency | 164360 |
| *ATP5F1B* | ATP synthase F1 subunit beta | 12 | Mitochondrial complex V: ATP synthase subunits | *atp5f1b* | 11 | *Atp5f1b* | 10 | -na- | -na- | 102910 |

| Approved symbol | Approved name | Chr | Gene family description | Zebrafish orthologs | Zebrafish chr | Mouse orthologs | Mouse chr | Clinical manifestations observed in humans | Disease | OMIM ID |
|---|---|---|---|---|---|---|---|---|---|---|
| *ATP5F1C* | ATP synthase F1 subunit gamma | 10 | Mitochondrial complex V: ATP synthase subunits | *atp5f1c* | 4 | *Atp5f1c* | 2 | -na- | -na- | 108729 |
| *ATP5F1D* | ATP synthase F1 subunit delta | 19 | Mitochondrial complex V: ATP synthase subunits | *atp5f1d* | 22 | *Atp5f1d* | 10 | Reduced assembly of complex V | Mitochondrial complex (ATP synthase) deficiency | 603150 |
| *ATP5F1E* | ATP synthase F1 subunit epsilon | 20 | Mitochondrial complex V: ATP synthase subunits | *atp5f1e* | 6 | *Atp5f1e* | 2 | Neonatal-onset lactic acidosis, 3-methylglutaconic aciduria, mild mental retardation, and peripheral neuropathy | Mitochondrial complex (ATP synthase) deficiency | 606153 |
| *ATP5IF1* | ATP synthase inhibitory factor subunit 1 | 1 | Mitochondrial complex V: ATP synthase subunits | *atp5if1a*<br>*atp5if1b* | 19<br>17 | *Atp5if1* | 4 | -na- | -na- | 614981 |
| *ATP5MC1* | ATP synthase membrane subunit c locus 1 | 17 | Mitochondrial complex V: ATP synthase subunits | *atp5mc1* | 3 | *Atp5mc1* | 11 | -na- | -na- | 603192 |
| *ATP5MC2* | ATP synthase membrane subunit c locus 2 | 12 | Mitochondrial complex V: ATP synthase subunits | -na- | -na- | *Atp5mc2* | 15 | -na- | -na- | 603193 |
| *ATP5MC3* | ATP synthase membrane subunit c locus 3 | 2 | Mitochondrial complex V: ATP synthase subunits | *atp5mc3a*<br>*atp5mc3b* | 9<br>6 | *Atp5mc3* | 2 | -na- | -na- | 602736 |
| *ATP5MD* | ATP synthase membrane subunit DAPIT | 10 | Mitochondrial complex V: ATP synthase subunits | *atp5md* | 1 | *Atp5md* | 19 | -na- | -na- | 615204 |
| *ATP5ME* | ATP synthase membrane subunit e | 4 | Mitochondrial complex V: ATP synthase subunits | *atp5mea*<br>*atp5meb* | 21<br>5 | *Atp5me* | 5 | -na- | -na- | 601519 |
| *ATP5MF* | ATP synthase membrane subunit f | 7 | Mitochondrial complex V: ATP synthase subunits | *atp5mf* | 3 | *Atp5mf* | 5 | -na- | -na- | -na- |
| *ATP5MG* | ATP synthase membrane subunit g | 11 | Mitochondrial complex V: ATP synthase subunits | *Atp5* | 5 | *Atp5mg* | 9 | -na- | -na- | 617473 |
| *ATP5MPL* | ATP synthase membrane subunit 6.8PL | 14 | Mitochondrial complex V: ATP synthase subunits | -na- | -na- | *Atp5mpl* | 12 | -na- | -na- | 604573 |
| *ATP5PB* | ATP synthase peripheral stalk-membrane subunit b | 1 | Mitochondrial complex V: ATP synthase subunits | *atp5pb* | 8 | *Atp5pb* | 3 | -na- | -na- | 603270 |
| *ATP5PD* | ATP synthase peripheral stalk subunit d | 17 | Mitochondrial complex V: ATP synthase subunits | *atp5pd* | 3 | *Atp5pd* | 11 | -na- | -na- | 618121 |
| *ATP5PF* | ATP synthase peripheral stalk subunit F6 | 21 | Mitochondrial complex V: ATP synthase subunits | *atp5pf* | 1 | *Atp5pf* | 16 | -na- | -na- | 603152 |
| *ATP5PO* | ATP synthase peripheral stalk subunit OSCP | 21 | Mitochondrial complex V: ATP synthase subunits | *atp5po* | 9 | *Atp5po* | 16 | -na- | -na- | 600828 |

| Approved symbol | Approved name | Chr | Gene family description | Zebrafish orthologs | Zebrafish chr | Mouse orthologs | Mouse chr | Clinical manifestations observed in humans | Disease | OMIM ID |
|---|---|---|---|---|---|---|---|---|---|---|
| *MT-ATP6* | mitochondrially encoded ATP synthase membrane subunit 6 | MT | Mitochondrial complex V: ATP synthase subunits | *mt-atp6* | MT | *mt-Atp6* | MT | Neurogenic muscle weakness, ataxia, and retinitis pigmentosa, lactic acidemia, hypotonia, neurodegenerative disease, mental retardation, infantile cardiac hypertrophy, sideroblastic anemia, delayed development, seizures and bilateral striatal necrosis | Leigh syndrome; Leber optic atrophy; Infantile mitochondrial bilateral striatal necrosis; Infantile hypertrophic cardiomyopathy; NARP syndrome | 516060 |
| *MT-ATP8* | mitochondrially encoded ATP synthase membrane subunit 8 | MT | Mitochondrial complex V: ATP synthase subunits | *mt-atp8* | MT | *mt-Atp8* | MT | Cardiac hypertrophy, neuropathy, dysarthric speech and ataxic gait | Hypertrophic cardiomyopathy | 516070 |
